# Loss of the actin remodeling protein Flightless-1 impairs CD8 and regulatory T cell function

**DOI:** 10.1101/2025.08.15.669900

**Authors:** Michelle M. Lissner, Jenna M. Sullivan, Minjian Ni, McKenna Sherve, Anne M. Hocking, Jessica A. Hamerman, Daniel J. Campbell

## Abstract

T cell immunity depends on the precise coordination of signaling networks with actin cytoskeleton remodeling, yet the molecular regulators of these processes remain incompletely defined. Flightless-1 (FLII) is a gelsolin-family actin regulator with unique leucine-rich repeats that can couple cytoskeletal dynamics to diverse signaling pathways. Here, using conditional knockout mice, we identify essential roles for FLII in both CD8^+^ and regulatory T cells. Loss of FLII in CD8^+^ T cells caused a profound loss of naive cells from the spleen, impaired CCR7-dependent migration, and defective accumulation in the lung parenchyma during antigen-specific responses to respiratory vesicular stomatitis virus infection, despite largely preserved activation, effector differentiation, and cytotoxic function. FLII-deficient Foxp3^+^ regulatory T cells maintained normal numbers but exhibited diminished CD25 expression, defective IL-2 signaling, and failed to restrain spontaneous, tissue-specific autoimmunity. These findings identify FLII as a critical and previously unrecognized orchestrator of T cell trafficking and immune regulation, which may link chemokine receptor signaling to actin remodeling and is essential for proper T cell migration and function.

## Introduction

T cell migration and stimulation are accompanied by dynamic changes in the actin cytoskeleton that are coordinated with activation of various signaling pathways (1). For instance, chemotactic migration of T cells relies on actin-mediated pseudopod protrusion at the leading edge of the cell, and propulsion by actin-dependent contractile forces in the trailing uropod. Additionally, T cell activation by antigen-presenting cells is accompanied by formation of a specialized cell-cell interface termed the ‘immunological synapse’ (IS), in which an actin ring helps constrain and localize essential transmembrane receptors and signaling components to ensure optimal TCR/co-stimulatory receptor signaling and activation (2). Formation of the IS also controls the directional release of cytotoxic effector molecules and cytokines from T cells that mediate target cell killing and/or activation. The importance of dynamic actin remodeling in T cells is highlighted by the large number of primary immunodeficiencies caused by loss of actin-regulatory proteins (3). These proteins are generally adaptors that help link activation of specific signaling pathways to coordinated changes in the actin cytoskeleton that control cell polarity, mobility, and activation.

Despite the clear importance of dynamic actin remodeling for proper T cell function, the molecular basis for how this is integrated with various signaling pathways is still poorly understood. A key protein family that regulates actin dynamics are proteins containing gelsolin domains, which function to regulate actin severing, branching, and capping (4). Among the gelsolin domain-containing proteins, Flightless-1 (FLII, encoded by the *Flii* gene) is highly conserved and has a unique domain structure, with its gelsolin domains linked to 16 N-terminal tandem leucine rich repeats (LRRs) that mediate interaction with a range of effector molecules that modulate different signaling pathways (5-10). Most relevant to T cell migration and activation, FLII binds to both RAC1 and the RAC1 GEF PREX1 during migration of fibroblasts (11), and can also directly interact with RAS and thereby regulate MAPK/ERK signaling (9). Thus, through its linked LRR and gelsolin domains, FLII is poised to integrate cell signaling with dynamic changes in the actin cytoskeleton to control CD8^+^ T cell migration and activation. Additionally, FLII has been implicated in control of gene expression, either by sequestration of key transcription/translation factors, or by directly acting as a transcriptional co-factor (12).

Despite its many potential functions, the role of FLII in controlling actin remodeling, signal transduction, and transcription during T cell migration and stimulation are almost completely unexplored. This is largely because FLII plays an essential role in gastrulation, and thus germline deletion of the *Flii* gene in mice is lethal at a very early stage in embryonic development (13). Although the function of FLII in T cells has been studied using *Flii*^*+/-*^ cells (14), these studies are difficult to interpret as this does not result in complete loss of FLII protein, and haploinsufficiency may not be sufficient to reveal most FLII functions. Therefore, to define the role of FLII in T cells, we have generated a floxed allele of the *Flii* gene that can be used to conditionally delete *Flii* in specific cell types. Crossing these animals to mice expressing Cre recombinase under control of the distal Lck promoter (dLck-Cre mice), we show that loss of FLII in CD8^+^ T cells impairs their homeostasis *in vivo*, reducing the overall number of cells and altering the balance of naïve and memory T cell subsets. FLII-deficient CD8^+^ T cells also showed altered responses and migration to the lungs following respiratory infection with vesicular stomatitis virus (VSV). The ability to form conjugates with antigen-presenting cells and undergo early events in T cell activation were not substantially altered in FLII-deficient cells, but their chemotactic responses to CCR7 ligands and their migration to the spleen were impaired. Furthermore, deletion of *Flii* in regulatory T (Treg) cells in Foxp3-Cre mice resulted in development of autoimmune inflammation in the skin and lungs associated with lymphadenopathy, splenomegaly and enhanced effector T cell differentiation. Thus, we identify FLII as a likely regulator of actin remodeling in T cells that is critical for normal T cell homeostasis and function.

## Results

### FLII is highly expressed in naïve and activated T cells

Because of the central role actin reorganization plays in T cell migration, activation and function, we leveraged a publicly-available dataset (GSE109125) to determine the expression of gelsolin-containing proteins in murine T cells (15). *Flii* was the most highly expressed family member in naïve CD4^+^ and CD8^+^ cells, followed by *Gsn* and *Svil* (**Fig 1A**). Examination of *Flii* and *Gsn* gene expression in T cell subsets further revealed that *Flii* gene expression is upregulated upon activation in both CD4 and CD8 cells, with highest expression in terminal effector and memory precursor CD8 T cells, whereas *Gsn* appeared to be downregulated upon T cell activation (**Fig 1B**). FLII is a large protein (predicted m.w. 143.6 kD) with a unique domain structure, with 16 tandem leucine rich repeats (LRRs) linked to 6 highly conserved C-terminal gelsolin domains (**Fig 1C**). To examine FLII protein levels in T cell subsets, we sorted CD44^lo^ naïve CD4^+^ and CD8^+^ T cells, and CD4^+^Foxp3^+^ regulatory T cells (T_R_) cells from the spleens of Foxp3-mRFP mice, and measured FLII protein levels by Western blot either directly after sorting, or after activation of the sorted naïve T cells with anti-CD3/CD28 for 48h (**Fig 1D**). FLII protein was detected in all T cell subsets, though at very low levels in naïve CD4^+^ T cells. Additionally, as suggested by analysis of *Flii* mRNA, cell activation increased FLII expression in both CD4^+^ and CD8^+^ T cells. Thus, FLII is expressed and poised to regulate cell signaling and actin remodeling in multiple T cell types.

**Figure 1:**
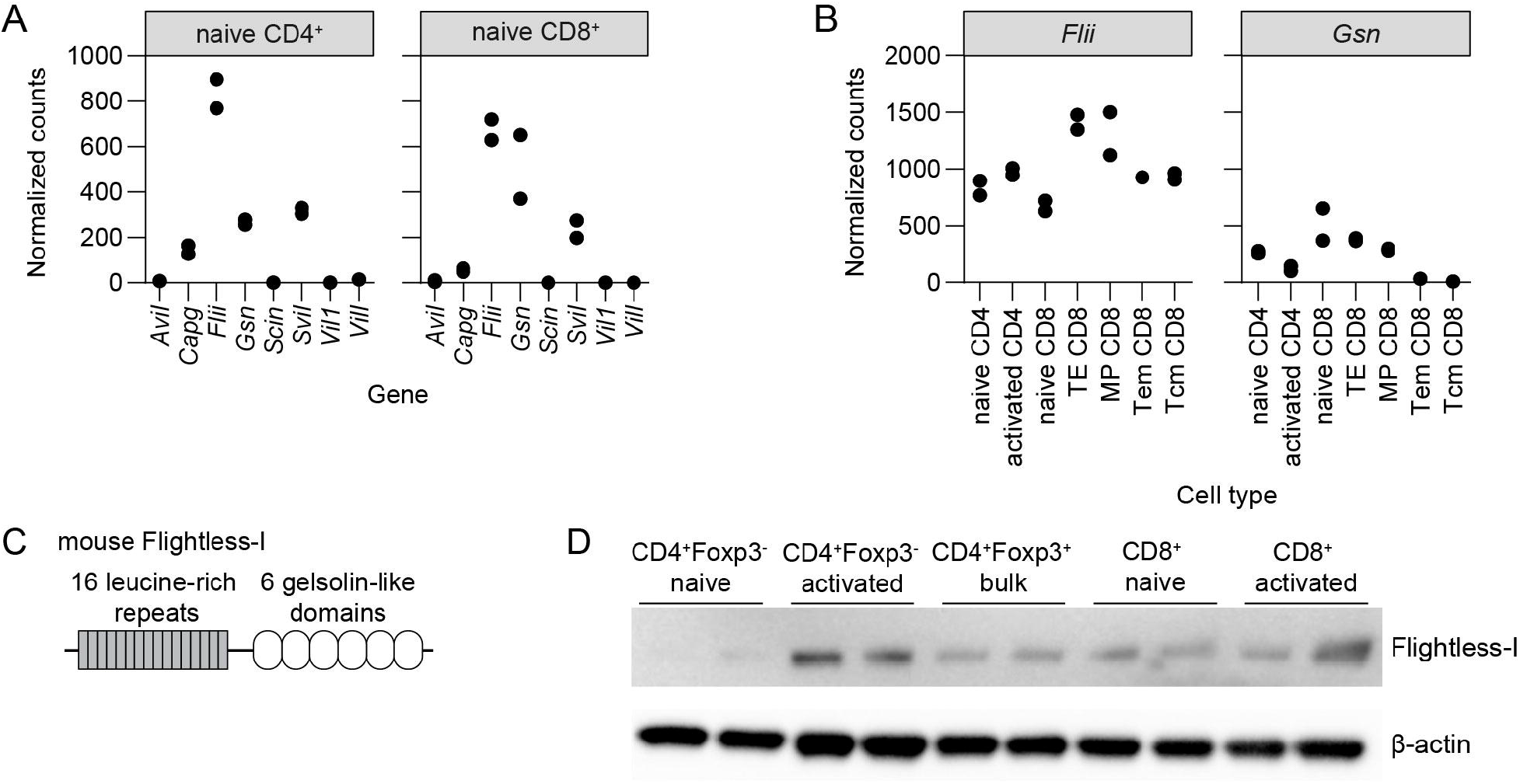
FLII is expressed in T cells. (A) Expression of the indicated genes gelsolin domains in murine naïve CD4^+^ and CD8^+^ T cells. (B) Expression of the *Flii* and *Gsn* genes in the indicated T cell populations. Data in (A) and (B) are from Gene Expression Omnibus dataset GSE109125. (C) Schematic of FLII protein structure. (D) Western blot analysis of FLII and β-actin protein expression in the indicated murine T cell populations. Data are representative of 2 independent experiments pooling cells from 2 mice per experiment.

### FLII is required for homeostasis of CD8^+^ T cells

Since germline knockout of the *Flii* gene is embryonic lethal, and studies of *Flii*^+/-^ T cells have been difficult to interpret (14), we generated a conditional knockout mouse model to study the role of FLII in T cell function. We crossed mice expressing a floxed allele of the *Flii* gene to mice expressing Cre recombinase in T cells under control of the distal Lck (dLck) promoter. Henceforth, we refer to these mice as *Flii*^*fl/fl*^*dLck*^*cre*^ mice and compared them to wild-type (*Flii*^*wt/wt*^*dLck*^*cre*^) animals. Western blot analysis of sorted splenocytes from *Flii*^*fl/fl*^*dLck*^*cre*^ mice showed a near-complete absence of FLII protein in CD8^+^ T cells, whereas FLII expression was maintained in both Foxp3^+^ and Foxp3^-^ CD4^+^ T cells (**Fig 2A**). This is consistent with prior reports of the *dLck*^*cre*^ allele functioning more efficiently in CD8^+^ cells (16), and therefore we focused our analyses of *Flii*^*fl/fl*^*dLck*^*cre*^ mice on potential alterations in the CD8^+^ T cell compartment. Importantly, we were able to study *Flii* deletion in mature T cells because the dLck promoter does not drive cre expression until after completion of positive selection in CD4^+^CD8^+^ ‘double positive’ thymocytes (17). Indeed, other than a small but significant decrease in the frequency of CD4^+^CD8^-^ mature single positive cells and a corresponding increase in the frequency of CD4^+^CD8^+^ double positive cells, thymic development appeared largely normal in *Flii*^*fl/fl*^*dLck*^*cre*^ mice (**Fig 2B**).

**Figure 2:**
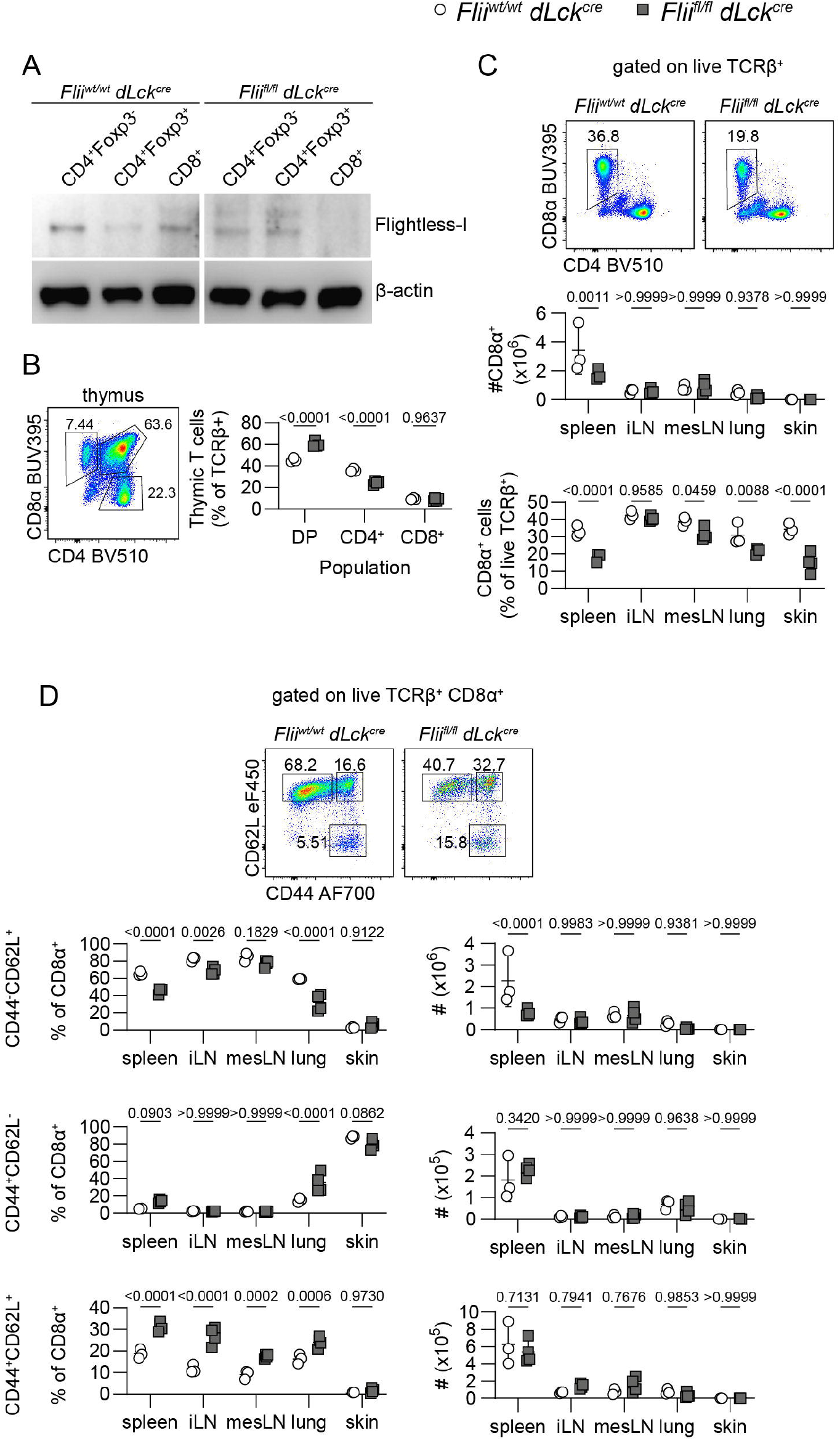
FLII deficiency alters CD8^+^ T cell frequency and phenotype *in vivo*. (A) Western blot analysis of FLII and β-actin protein expression in the indicated T cell subsets from sorted from *Flii*^*wt/wt*^*dLck*^*cre*^ and *Flii*^*fl/fl*^*dLck*^*cre*^ mice as indicated. (B) Representative flow cytometry staining of CD4 and CD8 expression by gated live thymocytes, and frequency of thymic T cell populations in *Flii*^*wt/wt*^*dLck*^*cre*^ and *Flii*^*fl/fl*^*dLck*^*cre*^ mice as indicated. (C) Representative flow cytometry staining of CD4 and CD8 by gated live TCRβ^+^ T cells in the spleens of *Flii*^*wt/wt*^*dLck*^*cre*^ and *Flii*^*fl/fl*^*dLck*^*cre*^ mice as indicated, and quantification of CD8^+^ cell number and frequency. (D) Representative flow cytometry staining and quantification of naïve (CD44^-^CD62L^+^), effector (CD44^+^CD62L^-^), and memory (CD44^+^CD62L^+^) CD8^+^ cells from *Flii*^*wt/wt*^*dLck*^*cre*^ and *Flii*^*fl/fl*^*dLck*^*cre*^ mice as indicated. Data are representative of 2 independent experiments, 2-4 mice per group. Statistical analyses performed by two-way ANOVA with Sidak’s multiple comparisons test.

*Flii* deletion did not impact total TCRβ^+^ cell number in the spleen, inguinal lymph node (iLN), mesenteric lymph node (mesLN), lung, or skin (**Fig S1A**). However, *Flii*^*fl/fl*^*dLck*^*cre*^ mice had decreased number and frequency of CD8^+^ cells in the spleen, as well as decreased frequency of CD8^+^ cells in the mesLN, lung, and skin, whereas there was no significant change in the frequency or number of CD8^+^ T cells in the iLN (**Fig 2C**). Among CD8^+^ T cells, the frequency of CD44^lo^CD62L^+^ naïve phenotype cells was dramatically reduced in the spleen, iLN, and lungs, with corresponding increases in the frequencies of CD44^hi^CD62L^+^ central memory (T_CM_) and CD44^hi^CD62L^-^ effector memory (T_EM_) phenotype cells (**Fig 2D, left**). Thus, although *Flii*^*fl/fl*^*dLck*^*cre*^ mice had a sharp decrease in the number of naïve phenotype CD8^+^ T cells in the spleen, the absolute numbers of CD44^hi^ effector/memory populations were not significantly changed in any tissue site (**Fig 2D, right**). In contrast to the defects we observed in CD8^+^ T cells, there was no significant effects on the absolute number or phenotype of either conventional CD4^+^Foxp3^-^ or regulatory CD4^+^Foxp3^+^ T cells in any tissue examined in *Flii*^*fl/fl*^*dLck*^*cre*^ mice, consistent with sustained FLII protein expression detected in both these population in these animals (**Fig S1 B-D**). Additionally, loss of FLII did not significantly affect IFN-γ production after stimulation with PMA/ionomycin (PMA/I) in either CD44^lo^ naïve or CD44^hi^ effector/memory CD8^+^ T cells (**Fig S1E**). Collectively these findings suggest that *Flii* is required for the proper homeostasis and maintenance of naïve CD8 T cells, particularly in the spleen.

### Loss of FLII alters the magnitude and localization of CD8^+^ T cell responses to viral infection

The altered abundance and phenotype of CD8^+^ T cells in *Flii*^*fl/fl*^*dLck*^*cre*^ mice led us to investigate how loss of FLII alters antigen-specific CD8^+^ T cell responses during viral infection *in vivo*. For this, we co-transferred equal numbers of congenically-marked ovalbumin (OVA)-specific TCR transgenic OT-1 cells from *Flii*^*wt/wt*^ (CD45.1/2^+^) and *Flii*^*fl/fl*^ (CD45.2^+^) *dLck*^*cre*^ mice into wild-type CD45.1^+^ recipients, which were infected with 10^4^ PFU of vesicular stomatitis virus (VSV)-OVA 24 hours later by intranasal administration (**Fig 3A-B**). We then compared the abundance, phenotype, and function of the transferred OT-1 cells in the spleen, mediastinal lymph node (medLN) and lungs at 7, 14, 21 and 28 days post-infection (dpi). In the spleens and lungs of infected animals, the frequency and number of FLII-deficient OT-1 cells were significantly lower than WT OT-1 cells at nearly all timepoints examined, whereas in the medLN the trend was similar but did not reach statistical significance other than at 7 dpi (**Fig 3C**). Within the lungs, we used intravenous labeling with anti-CD90.2-BV605 antibody to compare the localization of FLII-deficient and WT OT-1 cells in the tissue parenchyma (CD90.2^-^) and tissue vasculature (CD90.2^+^). The majority of transferred OT-1 cells at all timepoints examined were localized in the lung vasculature. However, compared to *Flii*^*wt/wt*^*dLck*^*cre*^ cells, *Flii*^*fl/fl*^*dLck*^*cre*^-derived cells showed impaired accumulation in the lung parenchyma at all timepoints examined (**Fig 3D**).

**Figure 3:**
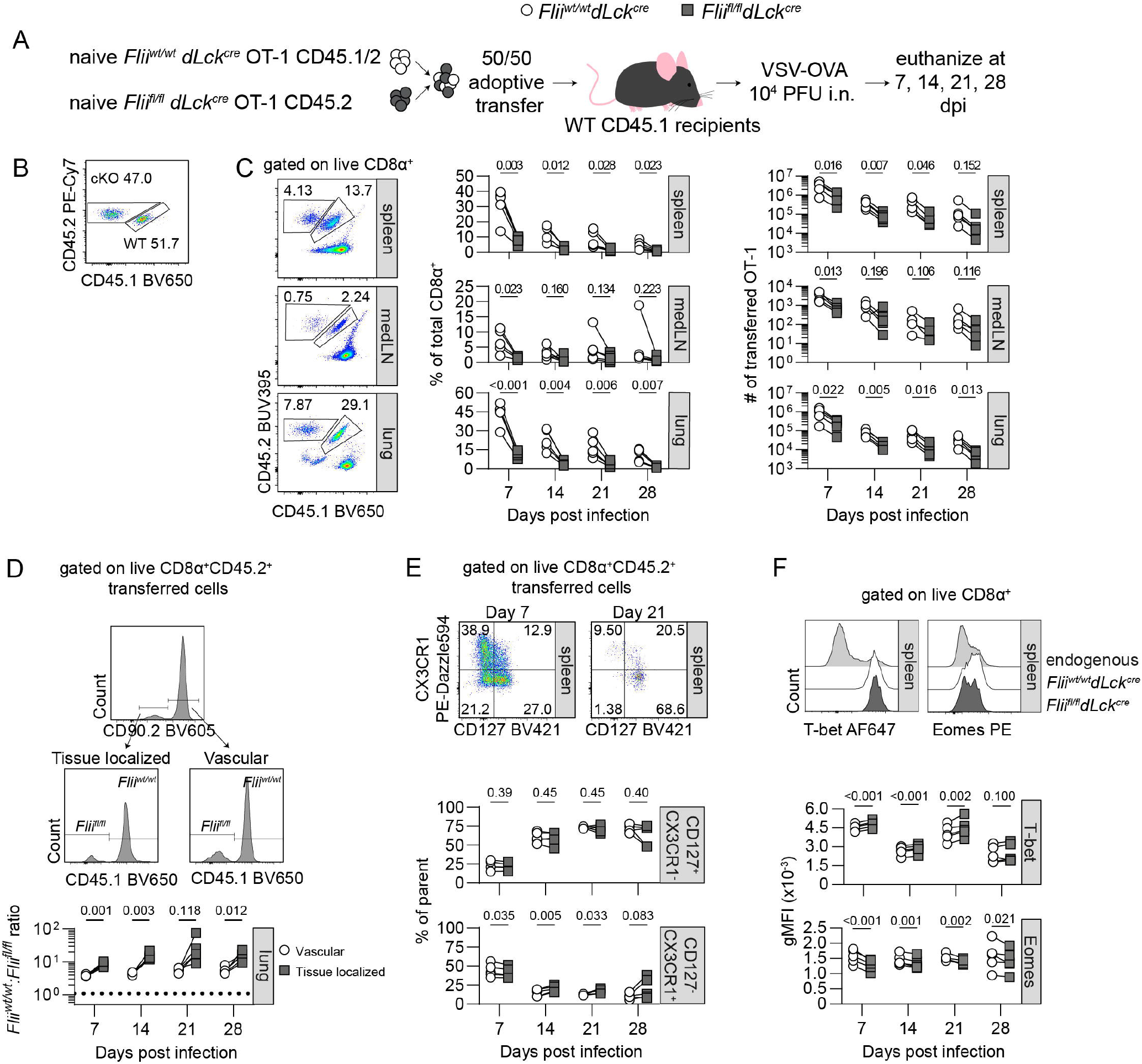
Altered response and localization of FLII-deficient CD8^+^ T cells during viral infection. (A) Experimental schematic. (B) Expression of CD45.2 and CD45.1 congenic markers by ingoing adoptively transferred OT-1 T cells. (C) Representative flow cytometry staining of CD45.2 and CD45.1 expression by gated live CD8^+^ T cells at 7 days post-infection in the indicated tissues, and quantification of the number and frequency of *Flii*^*wt/wt*^*dLck*^*cre*^ and *Flii*^*fl/fl*^*dLck*^*cre*^ OT-1 T cells in each tissue examined at different times post-infection as indicated. (D) Representative flow cytometry staining of CD90.2 and CD45.1 expression by gated live CD8^+^CD45.2^+^ transferred OT-1 T cells in the lungs at 7 days post-infection. Gates define intravascular CD90.2^+^ and parenchymal CD90.2^-^ populations. Quantification of the ratio of *Flii*^*wt/wt*^:*Flii*^*fl/fl*^ OT-1 T cells in each lung compartment at the indicated times post infection. (E) Representative flow cytometry staining of CX3CR1 and CD127 by gated CD8^+^CD45.2^+^ transferred OT-1 T cells in the spleens as the indicated times post-infection, and quantification of *Flii*^*wt/wt*^*dLck*^*cre*^ and *Flii*^*fl/fl*^*dLck*^*cre*^ OT-1 T cells with the indicated phenotype at each timepoint. (F) Representative flow cytometry staining of T-bet and Eomes expression by the indicated gated cell populations in the spleen at 7 days post infection, and quantification of Tbet and Eomes expression by *Flii*^*wt/wt*^*dLck*^*cre*^ and *Flii*^*fl/fl*^*dLck*^*cre*^ OT-1 T cells at each indicated timepoint. Data are representative of two independent experiments with 5 mice per group; connected points indicate data on WT and FLII-deficient T cells from the same mouse. Statistical analyses performed by paired T-test.

Despite the drop in cell number, loss of FLII had only minor impacts on the frequencies of CD8^+^ effector T cell subsets defined by differential expression of CD127 and CX3CR1 in the spleen (**Fig 3E**) or lungs (**Fig S2A**), although we did observe a small but significant shift in transcription factor expression with *Flii*^*fl/fl*^*dLck*^*cre*^ CD8^+^ cells expressing significantly increased levels of T-bet and decreased Eomesodermin (Eomes) in both the both tissue sites (**Fig 3F, Fig S2B**). To assess the functionality of FLII-deficient T cells, we compared production of the pro-inflammatory cytokines IFN-γ and TNF-α following ex vivo restimulation with either PMA/I or with the antigenic SIINFEKL peptide recognized by OT-1 T cells (**Fig S2C**, top and middle). Similarly, the ability of WT and FLII-deficient OT-1 T cells to release cytotoxic granules, as measured by surface translocation of the granule marker CD107 following stimulation, was comparable (**Fig S2C**, bottom). Thus, despite differences in the number and localization of virus-specific T cells, *Flii* deletion had minimal impacts on the differentiation and function of effector and memory CD8^+^ T cells.

### FLII regulates T cell migration

The decreased expansion of *Flii*^*fl/fl*^*dLck*^*cre*^ OT-1 T cells following VSV-OVA infection could be due to impaired activation and clonal expansion, or to a failure of the transferred naïve FLII-deficient OT-1 cells to adequately engraft at the site of viral T cell priming. Intranasal administration of VSV leads to widespread systemic infection and antigen-presentation in multiple secondary lymphoid tissues including the spleen (18). To examine the latter possibility, we determined the relative engraftment of naïve WT and FLII-deficient OT-1 T cells in different secondary lymphoid sites. We co-transferred equal numbers of ovalbumin-specific naive OT-1 cells from *Flii*^*wt/wt*^*dLck*^*cre*^ CD45.1/2^+^ and *Flii*^*fl/fl*^*dLck*^*cre*^ CD45.2^+^ donors into wild-type CD45.1^+^ recipient mice, and analyzed migration of these populations to the spleen, iLN, and mesLN 17h later (**Fig 4A, 4B**). Although there was no difference in the relative migration of WT and FLII-deficient OT-1 cells to either the iLN or mesLN, ∼3-fold fewer FLII-deficient T cells accumulated in the spleen relative to the co-transferred WT cells (**Fig 4C**). Although analysis of cell localization within the splenic red pulp vs. white pulp based on labeling with intravenous anti-CD90.2-BV605 antibody showed no statistically significant differences between WT and FLII-deficient cells, there was also a trend toward increased red pulp localization of FLII-deficient cells (**Fig 4D**).

**Figure 4:**
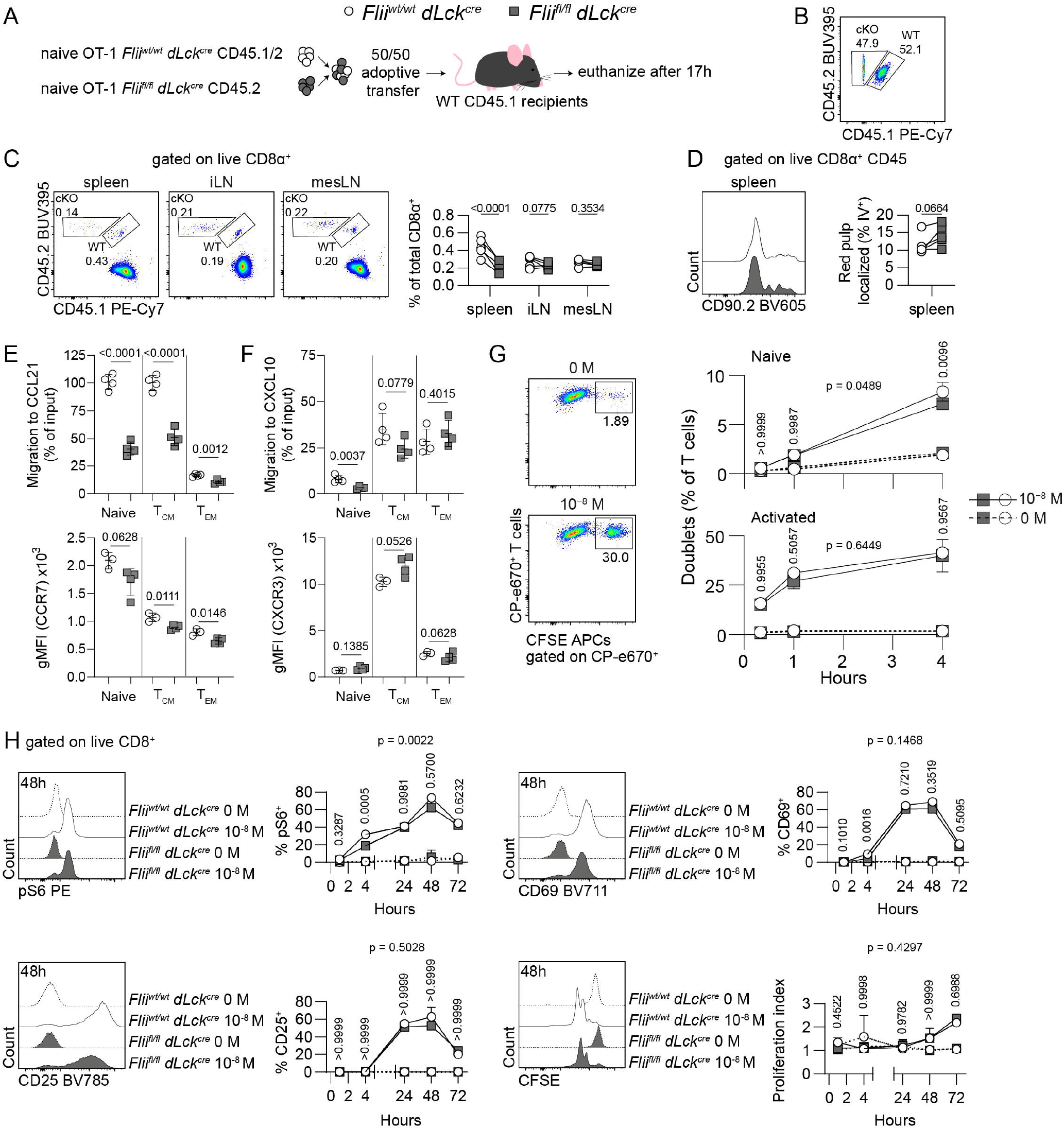
Loss of FLII Impairs CCR7-Directed Migration and Splenic Accumulation of Naïve CD8^+^ T Cells. (A) Experimental schematic. (B) Expression of CD45.2 and CD45.1 congenic markers by ingoing adoptively transferred OT-1 T cells. (C) Representative flow cytometry staining of CD45.2 and CD45.1 expression by gated live CD8^+^ T cells in each of the indicated tissues 24h after cell transfer, and quantification of the frequency of *Flii*^*wt/wt*^*dLck*^*cre*^ and *Flii*^*fl/fl*^*dLck*^*cre*^ OT-1 T cells in each tissue site. (D) Representative flow cytometry staining of CD90.2 expression by gated OT-1 T cells in the spleen, and quantitative analysis of the frequency of CD90.2+ red pulp localized cells of each genotype in the spleen. (E, F) Quantification of migration to CCL21 and expression of CCR7 (E), or migration to CXCL10 and expression of CXCR3 (F), by naïve, T_CM_ or T_EM_ splenic CD8^+^ T cell populations in *Flii*^*wt/wt*^*dLck*^*cre*^ and *Flii*^*fl/fl*^*dLck*^*cre*^ mice as indicated. (G) Representative flow cytometry staining of OT-1:APC doublets formed by preactivated OT-1 T cells from *Flii*^*wt/wt*^*dLck*^*cre*^ mice and APCs pre-cultured with or without SIINFEKL peptide as indicated, and quantification of the frequency of OT-1-APC doublets at the indicated timepoints and conditions. (H) Representative flow cytometry staining of pS6, CD25, CD69 and CFSE dilution by and OT-1 cells after *in vitro* stimulation with SIINFEKL-pulsed APCs, and quantitative analysis of marker expression at the indicated timepoints and conditions. *Data are representative of at least 2 independent experiments with 3-5 mice per group*. Statistical analyses by paired T-test (C-D); connected points indicate same mouse; by unpaired T-test (E-F); or by two-way ANOVA with Sidak’s multiple comparison test (G-H).

Migration of CD8^+^ T cells into secondary lymphoid tissues and inflamed tissue sites requires integrin activation and cellular chemotaxis mediated by chemokine receptor signaling. Thus, we examined expression and function of both CCR7 and CXCR3, key chemokine receptors that control migration of naïve and effector/memory CD8^+^ T cells, respectively. Interestingly, migration of FLII-deficient CD44^lo^CD62L^+^ naïve and CD44^hi^CD62L^+^ T_CM_ cells to the CCR7 ligand CCL21 was severely impaired relative to wild-type cells, and analysis of CCR7 expression by each of these populations showed modestly decreased CCR7 expression by FLII-deficient CD8^+^ T cells in all populations (**Fig 4E**). However, expression by FLII-deficient naïve T cells was still higher than in wild-type T_CM_, despite their much poorer migratory response. In contrast to the T cell responses to CCL21, migration to the CXCR3 ligand CXCL10 was not significantly impaired and expression of CXCR3 was normal or even slightly elevated in FLII-deficient T_CM_ and CD44^hi^CD62L^-^ T_EM_ (**Fig 4F**).

T cell activation is associated with extensive actin remodeling required for T cell conjugation to APCs, and is coordinated with cell signaling events that are essential for the clonal expansion and functional differentiation (2). Therefore, we determined if loss of FLII impairs T cell conjugation to peptide-loaded APCs, and alters T cell activation, signaling, and proliferation *in vitro*. To assess APC conjugation, CFSE-labeled splenocytes were loaded with 10^−8^M SIINFEKL peptide and incubated with naïve or pre-activated CP-e670-labeled OT-1 T cells from *Flii*^*wt/wt*^*dLck*^*cre*^ or *Flii*^*fl/fl*^*dLck*^*cre*^ mice for up to 4h, and the frequency of CFSE^+^CP-e670^+^ T cell-APC doublets was measured by flow cytometry. As expected, conjugate formation was not observed in the absence of SIINFEKL peptide and was much higher in OT-1 cells pre-activated with anti-CD3/CD28 compared with naïve OT-1 cells. Although these was no significant difference in conjugate formation between pre-activated wild-type and FLII-deficient cells, we did observe a small but significant impairment in the ability of FLII-deficient naïve T cells to form conjugates at the 4h timepoint (**Fig 4G**).

To determine how FLII affects T cell activation, we cocultured SIINFEKL-loaded splenocytes with OT-1 T cells and measured induction of T cell activation markers and cell proliferation by flow cytometry at various times between 4 and 72 hours (**Fig 4H**). Similar to what we observed for conjugate formation, TCR-mediated cell activation, as read out by phosphorylation of the S6 ribosomal protein (pS6) downstream of the PI3K/Akt/mTOR pathway, was mildly impaired in FLII-deficient T cells at 4h, but was normal at all other timepoints examined. Furthermore, upregulation of CD69 and CD25 were not altered by loss of FLII, and neither the kinetics nor magnitude of proliferation as measured by CFSE dilution were impacted by FLII deficiency. Together, these data indicate that FLII plays only a minor role in the early events in cell interaction, immune synapse formation, or cell signaling/activation. Instead we favor a model in which FLII is required for migration and localization within specific tissue environments such as the spleen and lung parenchyma, and that this results in impaired homeostasis and activation of FLII-deficient CD8^+^ T cells.

### FLII is required for regulatory T cell function in vivo

To determine if FLII regulates the abundance and activity of other T cell populations, we generated *Flii*^*fl/fl*^*Foxp3*^*YFPcre/Y*^ mice to selectively delete FLII expression in Treg cells. As proper Treg activity is required to maintain immune homeostasis and prevent catastrophic autoimmunity (19, 20), this is a sensitive system to assess protein function in T cells. Although young (5-7 week-old) *Flii*^*fl/fl*^*Foxp3*^*YFPcre/Y*^ mice appeared healthy, they tended to weigh less than age-matched controls, and stopped gaining weight around 8-10 weeks of age (**Fig 5A**). Additionally, *Flii*^*fl/fl*^*Foxp3*^*YFPcre/Y*^ mice developed spontaneous dermatitis on the tail and ears by 13-15 weeks of age (**Fig 5B**) and had evident lymphadenopathy and splenomegaly indicative of immune dysregulation (**Fig 5C**). Histological analysis confirmed inflammatory dermatitis in *Flii*^*fl/fl*^*Foxp3*^*YFPcre/Y*^ mice, and we also found inflammatory pathology in the lungs of these animals (**Fig 5D**, left and middle). However the liver, a major target organ of inflammation in Foxp3-deficient mice (20), appeared histologically normal (**Fig 5D**, right). In 13-15 week-old *Flii*^*fl/fl*^*Foxp3*^*YFPcre/Y*^ mice, we observed decreased Treg cell frequency in the mesLN and skin, but the total number of Treg cells present in each tissue examined was not significantly different than in *Flii*^*wt/wt*^*Foxp3*^*YFPcre/Y*^ control animals (**Fig 5E**). However, in most tissues examined, *Flii*^*fl/fl*^*Foxp3*^*YFPcre/Y*^ mice had a higher fraction of CD44^+^CD62L^-^ effector Treg cells than controls (**Fig 5F**) (21), along with other features of activated Treg cells including decreased expression of CD25, elevated expression of the proliferation marker Ki-67, and higher expression of GITR that is upregulated upon Treg cell activation (**Fig S3A**) (22). Decreased CD25 expression was associated with diminished IL-2 signaling in Treg cells as measured by STAT5 phosphorylation directly ex vivo in both the spleen and iLN (**Fig S3B**). Indicative of impaired Treg cell function and an inability to properly restrain T cell activation, both CD4^+^Foxp3^-^ Tconv and CD8^+^ cells were significantly more activated in *Flii*^*fl/fl*^*Foxp3*^*YFPcre/Y*^ mice than in controls (**Fig 5G, Fig S4A**). Additionally, IFN-γ and IL-17A production were increased in splenic CD4^+^ Tconv from *Flii*^*fl/fl*^*Foxp3*^*YFPcre/Y*^ mice, and a small fraction of both Treg cells and CD8^+^ T cells also acquired the ability to produce IL-17A, whereas TNF-α production was decreased in both Treg and Tconv cells (**Fig S4B**). To determine if the altered Treg cell phenotypes we observed in *Flii*^*fl/fl*^*Foxp3*^*YFPcre/Y*^ mice were cell intrinsic or secondary to the inflammatory phenotype that develops in these animals, we examined *Flii*^*fl/fl*^*Foxp3*^*YFPcre/mRFP*^ or *Flii*^*wt/wt*^*Foxp3*^*YFPcre/mRFP*^ heterozygous female mice, in which the presence of FLII-sufficient Treg cells due to random X-chromosome inactivation prevents development of inflammatory disease. CD25 expression remained significantly lower in YFP^+^ Treg cells from *Flii*^*fl/fl*^*Foxp3*^*YFPcre/mRFP*^ compared with those from *Flii*^*wt/wt*^*Foxp3*^*YFPcre/mRFP*^ animals (**Fig S4C, left**). However, enhanced effector Treg cell differentiation indicated by elevated expression of CD44 was not observed in the FLII-deficient Treg cells from *Flii*^*fl/fl*^*Foxp3*^*YFPcre/mRFP*^ mice, and thus this phenotype is likely secondary to the inflammatory disease that develops in the *Flii*^*fl/fl*^*Foxp3*^*YFPcre/Y*^ animals. (**Fig S4C, right**). Together, these data demonstrate that in addition to a functional role in CD8^+^ T cells, FLII is essential for proper Treg activity, suggesting a broader role for this protein in the regulation of T cell function.

**Figure 5:**
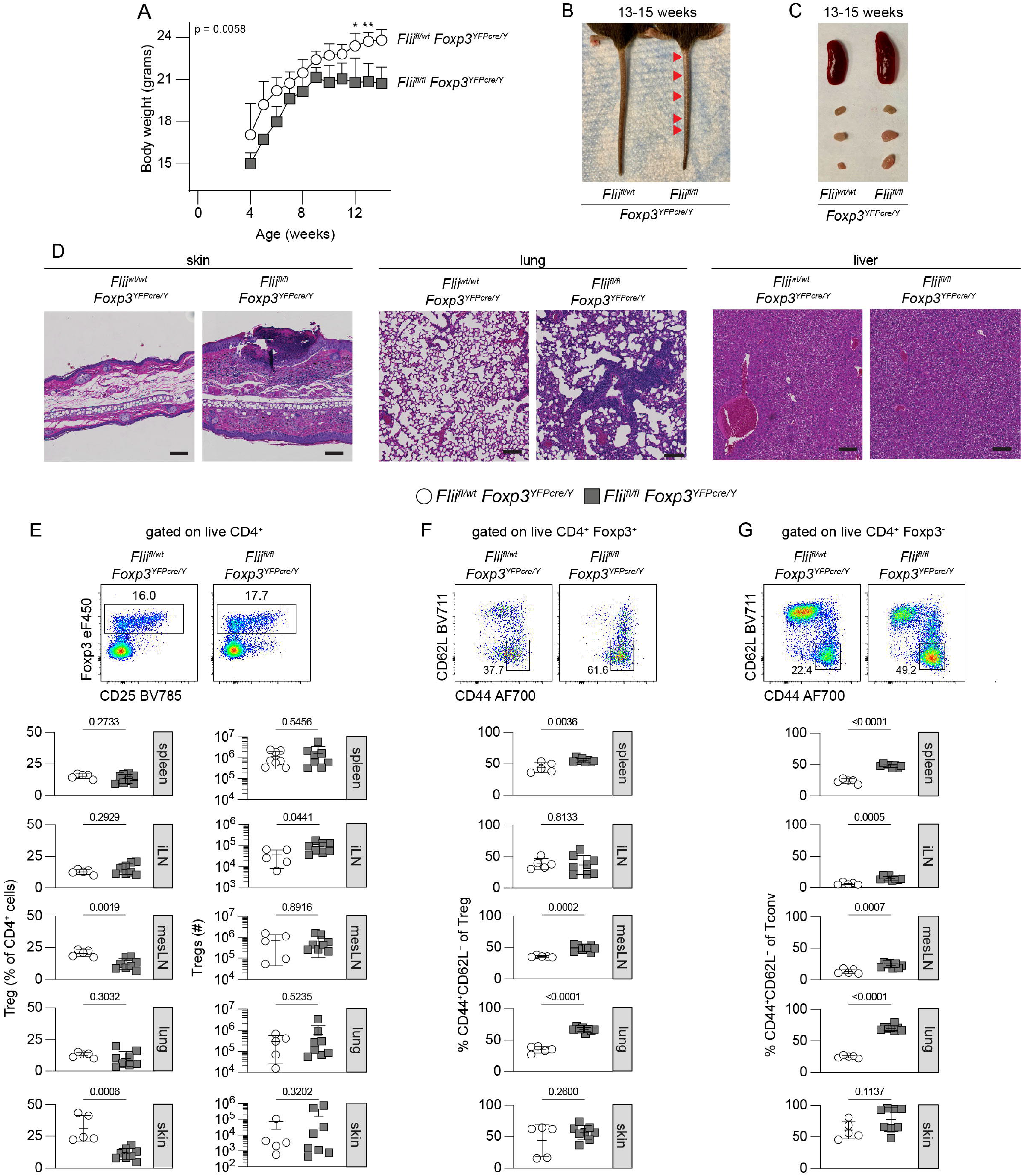
Spontaneous inflammatory disease in mice lacking FLII in Treg cells. (A) Body weight curve of *Flii*^*fl/fl*^*Foxp3*^*YFPcre/Y*^ and *Flii*^*wt/wt*^*Foxp3*^*YFPcre/Y*^ mice as indicated. (B) Representative images of tail skin of *Flii*^*fl/fl*^*Foxp3*^*YFPcre/Y*^ and *Flii*^*wt/wt*^*Foxp3*^*YFPcre/Y*^ mice as indicated at 13-15 weeks of age. (C) Representative images of spleen and peripheral lymph nodes (inguinal, axillary and brachial) of *Flii*^*fl/fl*^*Foxp3*^*YFPcre/Y*^ and *Flii*^*wt/wt*^*Foxp3*^*YFPcre/Y*^ mice as indicated at 13-15 weeks of age. (D) H&E staining of skin, lung, and liver sections from *Flii*^*fl/fl*^*Foxp3*^*YFPcre/Y*^ and *Flii*^*wt/wt*^*Foxp3*^*YFPcre/Y*^ mice as indicated at 13-15 weeks of age. Scale bar = 100μM. (E) Representative flow cytometry staining of Foxp3 and CD25 by gated live CD4^+^ T cells from *Flii*^*fl/fl*^*Foxp3*^*YFPcre/Y*^ and *Flii*^*wt/wt*^*Foxp3*^*YFPcre/Y*^ mice as indicated, and quantification of Treg cell frequency and number from the indicated tissues at 13-15 weeks of age. (F) Representative flow cytometry staining of CD44 and CD62L expression by gated live CD4^+^Foxp3^+^ Treg cells from *Flii*^*fl/fl*^*Foxp3*^*YFPcre/Y*^ and *Flii*^*wt/wt*^*Foxp3*^*YFPcre/Y*^ mice as indicated, and quantification of the frequency of CD44^+^CD62L^-^ effector Treg cells from the indicated tissues at 13-15 weeks of age. (G) Representative flow cytometry staining of CD44 and CD62L expression by gated live CD4^+^Foxp3^-^ Tconv cells from *Flii*^*fl/fl*^*Foxp3*^*YFPcre/Y*^ and *Flii*^*wt/wt*^*Foxp3*^*YFPcre/Y*^ mice as indicated, and quantification of the frequency of CD44^+^CD62L^-^ activated T cells from the indicated tissues at 13-15 weeks of age. Data indicate 3-8 mice per condition combined from at least 5 independent experiments. Statistical analyses by two-way ANOVA with Sidak’s multiple comparisons test.

## Discussion

The integration of cell signals and cytoskeletal remodeling is essential for the dynamic functions of T cells as they migrate through lymphoid and non-lymphoid tissues, recognize their cognate antigens and undergo robust clonal expansion, and carry out both pro- and anti-inflammatory effector responses. We have made the novel observation that loss of FLII protein expression results in impaired peripheral maintenance and clonal expansion of CD8^+^ T cells and altered migratory responses in vitro and in vivo. We further showed that loss of FLII expression in Foxp3^+^ regulatory T cells causes progressive loss of tolerance, accumulation of effector phenotype CD4^+^ and CD8^+^ T cells in multiple tissues, and inflammatory disease in the skin and lungs. Based on its unique domain structure, we hypothesize that FLII links changes in the actin cytoskeleton to the activation and spatial organization of key signaling pathways during T cell migration and activation, thereby acting as an important regulator of T cell responses.

The critical function of cytoskeletal remodeling proteins in T cell function is evidenced by the large number of primary immune deficiencies caused by hypomorphic or non-functional mutations in proteins that control this process. These include genes such as *WAS, DOCK2, DOCK8, CORO1A, and ARPC1B* (23-27). These proteins all act as integrators that coordinate cytoskeletal remodeling in response to activation of specific signaling pathways. Although no primary immune deficiencies have been associated with mutations in the *Flii* gene, this may be due to the fact that unlike DOCK2, DOCK8 and WASP whose expression is largely limited to immune cells, *Flii* is expressed in nearly all cells and germline knockout of *Flii* in mice is embryonic lethal (13). Our use of a novel floxed *Flii* allele in combination with two Cre-expressing lines that efficiently delete *Flii* in either CD8^+^ T cells or CD4^+^Foxp3^+^ regulatory T cells reveals an essential function for this protein in controlling the responses of these cells.

In CD8^+^ T cells, loss of FLII expression led to a complex phenotype characterized by diminished CD8^+^ T cell frequencies and numbers in multiple tissue sites, especially the spleen. This was particularly associated with a substantial loss of CD44^lo^CD62L^hi^ naïve phenotype CD8^+^ T cells, and a corresponding increased frequency of CD44^hi^ effector/memory phenotype cells. The reason for the dramatic loss of naïve CD8^+^ T cells is not clear, but may be linked to decreased expression and function of CCR7 in FLII-deficient cells. Fibroblastic reticular cells (FRCs) in the secondary lymphoid tissues express high levels of the CCR7 ligands CCL19 and CCL21, and are the primary source of the cytokine IL-7 that promotes the survival and homeostasis of naïve CD4^+^ and CD8^+^ T cells (28). Although FLII-deficient naïve T cells migrated to LNs as well as WT cells following adoptive transfer, their localization within lymph nodes may be altered, resulting in impaired homeostasis. Interestingly, FLII*-*deficient naïve T cells showed defective accumulation in the spleen upon adoptive transfer, and the number of naïve splenic CD8^+^ T cells was selectively impacted by loss of FLII. Naive T cell trafficking to the lymph nodes requires activation of the LFA-1 integrin by CCR7 signaling triggered as cells role along high endothelial venules (HEV). By contrast there are no HEV in the spleen, and T cells are retained in the spleen and migrate into the splenic T cells zones following chemotactic gradients via CRR7 through white pulp ‘bridging channels’ (29). Thus, the selective failure of FLII-deficient T cells to accumulate is consistent with their impaired chemotaxis in response to CCL21. Interestingly, the ability of FLII-deficient memory T cells to migrate to the CXCR3 ligand CXCL10 was not altered, suggesting FLII has a specific function in CCR7-dependent T cell migration. Additionally, the selective loss of naïve CD8^+^ T cells from the spleen is similar to what is found in *Dock8*^*-/-*^ CD8^+^ T cells (30), suggesting that FLII and DOCK8 may both function in a common process that supports CD8^+^ T cell migration and homeostasis. Among its interaction partners, FLII directly binds to the signaling adaptor BCAP (encoded by the *Pik3ap1* gene), and this complex inhibits activation of the NLRP3 and NLRC4 inflammasomes in macrophages (10). BCAP also regulates CD8^+^ T cell differentiation and survival after activation (31, 32). However, unlike FLII BCAP is not expressed in naïve T cells, and the phenotypes of CD8^+^ T cells lacking BCAP or FLII are distinct, with BCAP-deficient cells showing impaired survival of effector T cells after clonal expansion, associated with decreased expression of T-bet and KLRG1 and increased Eomes expression. Thus we posit that most key functions of FLII in CD8^+^ T cells are either BCAP-independent, or that FLII and BCAP may antagonize each other function to fine tune CD8^+^ T cell differentiation.

Despite the defects in naïve CD8^+^ T cell homeostasis and migration, we found that the ability of FLII-deficient OT-1 T cells to expand and compete with WT cells in an anti-viral T cell response was only mildly impaired. Although the percentage and number of FLII-deficient cells was decreased ∼3x compared to WT cells at all timepoints and tissues examined after infection, this is similar to the defect in migration of naïve OT-1 T cells to the spleen after adoptive transfer. The spleen is a major site of infection and T cell priming following intranasal VSV infection, and thus the decreased number of FLII-deficient T cells likely reflects this initial seeding rather than impaired T cell activation or proliferation. Indeed, both naïve and previously activated FLII-deficient OT-1 T cells formed conjugates with peptide-pulsed APCs equivalently to WT cells, and early TCR signaling and activation as read out by phosphorylation of S6, induction of the activation markers CD25 and CD69, and cell proliferation were all similar in WT and FLII-deficient T cells. The functional differentiation of FLII-deficient T cells was only slightly altered relative to WT cells, further indicating that FLII does not appear to have a major role in modulating CD8^+^ T cell activation, clonal expansion and function.

Consistent with a broader function for FLII in regulating T cell activity, we found that deletion of *Flii* selectively in Foxp3^+^ cells led to Treg dysfunction as indicated by development of spontaneous immune dysregulation and tissue-specific inflammatory disease. Interestingly, unlike FLII-deficient CD8^+^ T cells, Treg cell numbers were normal in all tissues examined in *Flii*^*fl/fl*^*Foxp3*^*YFPcre/Y*^ mice, including the spleen, suggesting that FLII may regulate the localization and function of CD8^+^ T cells and Treg cells by distinct mechanisms. Similar to what we observed with CD8^+^ T cells, Treg cells showed increased activation, with fewer CD44^lo^CD62L^+^ central Treg cells in *Flii*^*fl/fl*^*Foxp3*^*YFPcre/Y*^ mice. However, accumulation of effector phenotype Treg cells is a common feature in inflammatory disease caused by Treg dysfunction, and analysis of FLII-deficient Treg cells in *Foxp3*^*YFPcre/mRFP*^ heterozygous mice that do not develop autoimmunity showed that these phenotypes were likely secondary to this inflammation, and not a direct consequence of FLII deficiency. However, decreased CD25 expression was a common feature in both *Flii*^*fl/fl*^*Foxp3*^*YFPcre/Y*^ and *Flii*^*fl/f*^*Foxp3*^*YFPcre/mRFP*^ heterozygous mice, and diminished responsiveness to IL-2 may underlie the Treg cell dysfunction we observed. This is similar to DOCK8-deficient Treg cells, which show impaired CD25 expression and decreased IL-2 responsiveness (33). The similarity in both CD8^+^ T cell and Treg cell phenotypes in FLII- and DOCK8-deficient mice suggest these proteins may function in some common pathways to help coordinate T cell migration and activation. However, in contrast to the phenotypes we observed, mice lacking DOCK8 expression in Treg cells also show decreased Treg abundance, and develop severe intestinal inflammation whereas skin was not affected (33), indicating non-overlapping functions for these proteins in T cell activation and migration to specific tissue sites.

Together, our findings identify FLII as a novel and essential regulator of T cell homeostasis, trafficking, and immune regulation. Given its widespread expression and its ability to integrate cytoskeletal remodeling with cell signaling networks, further investigation into the molecular interactions and downstream pathways regulated by FLII in T cells may uncover new mechanisms linking cell activation to actin remodeling to control cell migration and homeostasis.

## Materials and Methods

### Mice

All mice were maintained at Benaroya Research Institute, and experiments were preapproved by the Institutional Animal Care and Use Committee of Benaroya Research Institute. C57BL/6J, B6-SJL, OT-1, dLck-cre, Foxp3-YFPcre, and Foxp3-mRFP mice were purchased from The Jackson Laboratory. C57BL/6N-*Flii*^tm1a^ mice were generated by KOMP and purchased from the Mouse Biology Program at University of California Davis (Stock Number: 055670-UCD). To remove the inserted Frt-flanked lacZ/neo cassette, these mice were crossed to the FLP recombinase expressing strain B6.Cg-Tg(ACTFLPe)9205Dym/J (Jackson Laboratory) resulting in *Flii*^*fl/wt*^ mice with loxP sites flanking exons 8-14. Mice used in experiments were between 8-12 weeks of age at the time of sacrifice unless otherwise indicated.

### Intravascular labeling

Mice were anesthetized with 4% isoflurane and injected retroorbitally with 3μg of BV605 Thy1.2 (Biolegend) in 200 μl sterile PBS (Cytiva) 3 min before sacrifice. Tissue digestion and flow cytometry was performed as described below and flow cytometry used to define vascular (BV605^+^) and tissue-localized (BV605^-^) cells.

### Tissue digestion

Whole spleens and lymph nodes were mechanically disrupted and passed through 70μm strainers (Fisher) into RPMI 1640 (Cytiva) with 10% FBS (Sigma) (R10). Splenic erythrocytes were lysed using ACK lysis buffer (Gibco) and the remaining cells were resuspended in R10. Minced lungs were incubated in 5 ml RPMI 1640 with 50μg/ml Liberase™ (Roche) and 10U/ml DNAse I (Sigma) for 45 min while shaking at 200 rpm at 37°C. Digestion was quenched with 5 ml R10 and tissue was passed through a 70μm filter. Erythrocytes were lysed by ACK lysis buffer and the remaining cells were resuspended in R10. Ear skin was peeled into two layers and rinsed in R10 before being minced and incubated in RPMI 1640 with 27μl of 26U/mL Liberase™ and 25μl of 20mg/ml DNase I. Tissues were shaken at 200 rpm at 37°C. After 45 minutes, digestion was quenched with 5ml R10 and tissues were passed through a 70μm strainer. Cells were resuspended in R10.

### Flow cytometry

Single cell suspensions were prepared as described above. Cells were stained for surface antigens for 20 min in PBS at 4°C, fixed and permeabilized with Foxp3/Transcription Factor Staining Buffer Set (ThermoFisher) for 30 min at 4°C, and stained for intracellular antigens at RT for 30 minutes. Unless otherwise noted, surface antibody stains were performed at 4°C for 20 min and intracellular stains were performed at RT for 30 minutes. To measure CCR7, single cell suspensions were rested at 37°C, 5% CO_2_ in complete medium for 1 hour before incubating with CCL19-human IgG-Fc fusion protein (34). Cells were washed with 0.5% BSA (Sigma) in PBS (Cytivia) and incubated with goat anti-human Fc-PE (Jackson Immunoresearch). To measure pSTAT5, approximately one quarter of each spleen or one entire lymph node was mechanically disrupted using glass slides into Fixation/Permeabilization Buffer from Foxp3/Transcription Factor Staining Buffer Set (eBioscience) immediately upon euthanasia. Cells were incubated on ice, washed in FACS buffer, and resuspended in ice-cold 90% methanol (Fisher) for at least 30 min; cells were then washed with Fixation/Permeabilization Buffer and stained for flow cytometry for 45 min at room temperature. To measure ex vivo cytokine production, single-cell splenocyte suspensions were treated with 50 ng/ml PMA (Sigma), 1 μg/ml ionomycin (Sigma), and 2 μM monensin (eBioscience) in R10 for 4 hours at 37°C, 5% CO_2_ before staining. Alternatively, splenocytes were incubated with 10^−8^nM SIINFEKL peptide (InvivoGen) in R10 for 2 hours before adding 2 μM monensin and incubation for 4 more hours at 37°C, 5% CO_2_ before staining. Counting beads (Polysciences) were used to enumerate cells. Flow cytometric data were acquired on BD Symphony II flow cytometers (BD Biosciences) and data were analyzed using FlowJo software. In some experiments, naïve CD8^+^ T cells were isolated by positive selection anti-CD8a microbeads (Miltenyi), and subsequent sorting (live CD44^-^ CD62L^+^ CD8a^+^ TCR Vα2^+^ TCR Vβ5^+^) on a FACS Aria Fusion or FACS Aria III (BD).

### Adoptive transfer and infection

OT-1 CD8 T cells were isolated from *Flii*^*fl/fl*^*dLck*^*cre*^ or *Flii*^*wt/wt*^*dLck*^*cre*^ as described above. For experiments without subsequent infection, 5×10^5^ cells of each genotype were co-transferred into congenically-marked recipient mice via retroorbital intravenous injection. Tissues were harvested and processed at the indicated timepoints as described above. For experiments with subsequent infection 5×10^3^ cells of each genotype were co-transferred into congenically-marked recipient mice via intravenous injection one day before intranasal infection with 10^4^ PFU of VSV-OVA (35). Tissues were harvested and processed at the indicated timepoints as described above.

### T cell-APC conjugation

OT-1 CD8 T cells were isolated as described above and labeled with 1μM CP-e670 (ThermoFisher) for 9 minutes. Wild-type splenocytes were labeled with 1μM CFSE (Life Technologies) for 9 minutes, incubated with 0 M or 10^−8^ M SIINFEKL peptide for 1h at 37°C, and washed. An equal number of T cells and APCs (1-5×10^5^) were cocultured for the indicated timepoints. To measure conjugation, cells were fixed with PFA (Sigma) at the indicated timepoints and conjugate formation was read out by flow cytometry as % CFSE^+^ events of total CP-e670^+^ events.

### T cell stimulation and proliferation assays

OT-1 splenocytes were labeled with 1μM CFSE for 9 minutes. Wild-type splenocytes were incubated with 0 M or 10^−8^ M SIINFEKL peptide for 1h at 37°C and washed. 1×10^5^ T cells and 1×10^5^ APCs were cocultured for the indicated timepoints before reading our cell activation (by expression of pS6, CD25 and CD69) or proliferation (by dilution of CFSE) by flow cytometry.

### Western blots

CD8^+^ cells were isolated by MACS as described above. CD4^+^ cells were isolated by MACS using CD4 L3T4 microbeads (Miltenyi), and sorted in Foxp3^+^ and Foxp3^-^ fractions. Cells were resuspended in lysis buffer consisting of RIPA buffer (50 mM Tris–HCl (pH 7.4), 150 mM NaCl, 1% Triton X-100, 0.5% sodium deoxycholate, 0.1% SDS, and 5 mM EDTA in water) with protease inhibitors (Roche) and 1nM sodium orthovanadate (Sigma) and incubated on ice for 20 min. Lysates were spun at max speed for 10 min at 4°C before supernatants were collected. Protein concentration was determined by Pierce BCA protein assay (Thermo Scientific) before normalization and the addition of 1 part Laemmli Sample Buffer (Bio-Rad) to 3 parts lysate. Lysate was loaded onto a NuPAGE 4 to 12% Bis-Tris Gel, 1.5 mm x 15 wells (Invitrogen) after mixing with 1X NuPAGE LDS Sample Buffer (Invitrogen), and run at 120 V for 2.5 h. Samples were transferred onto a PVDF membrane (Millipore) at 4°C at 250 mA for 1.5 h before blocking in PBS with 1% Tween (Sigma) and 5% BSA. Membranes were stained with primary antibodies against FLII (Santa Cruz) and β-actin (Cell Signaling Technology) and secondary antibodies anti-mouse IgG HRP (Invitrogen) and anti-rabbit IgG HRP (Cell Signaling Technology) before imaging on a Chemidoc (BioRad) using Immobilon Western Chemiluminescent HRP Substrate (Millipore).

### Ex vivo migration assay

Single cell suspensions were from indicated tissues were rested for 1 hour at 37°C/5% CO_2_ to allow cytokine receptor levels to equilibrate. 100nM CXCL10 (Peprotech) or CCL21 (Peprotech) were plated in the bottom compartment of 24-well transwell plates (Corning Costar) and 1×10^6^ cells were plated in the transwell inserts. After 90 minutes of incubation at 37°C and 5% CO_2_, transwell inserts were discarded and migrated cells were stained for flow cytometry analysis and enumerated using counting beads.

### Histology

Tissues were excised and immediately fixed in 10% formalin for 24 hours before tissues were paraffin embedded. H&E staining was performed on 6-µm tissue sections by the Benaroya Research Institute Histology Core. Full slide scans were performed with the ImageXpress Micro Confocal High-Content Imaging System (Molecular Devices).

### Statistical analysis

All data are presented as mean values +/-SD and graphs were generated and analyzed using Prism software (GraphPad). Statistical analyses were performed as described in the figure legends.

## Acknowledgements

We would like to thank Pam Johnson and Caroline Stefani for assistance with histology and imaging; Adam Wojno and Adin Pierce for help with flow cytometry; Andrew Burch and the BRI veterinary staff for support with mouse work. This work was supported by grants R01AI124693 and R21AI172140 and R21AI157440 from the NIH to D.J.C., and grants R01AI150178 and R01AI113325 to JAH. MML was supported by a training grant from the NIH to the University of Washington (T32AR007108).

