## Supplementary figures and images for "Loss of the actin remodeling protein Flightless-1 impairs CD8 and regulatory T cell function"

### Fig S1

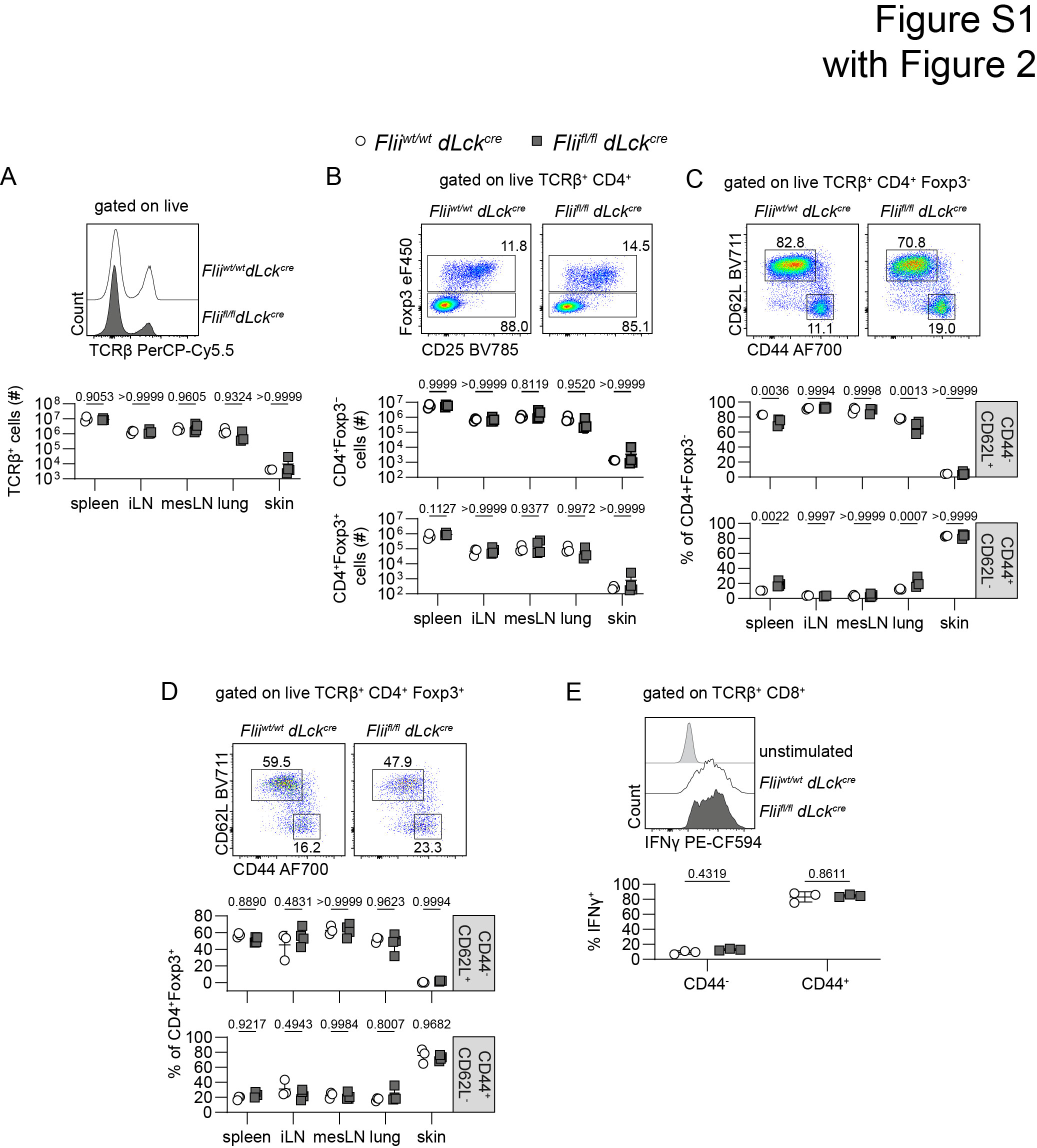

### Fig S2

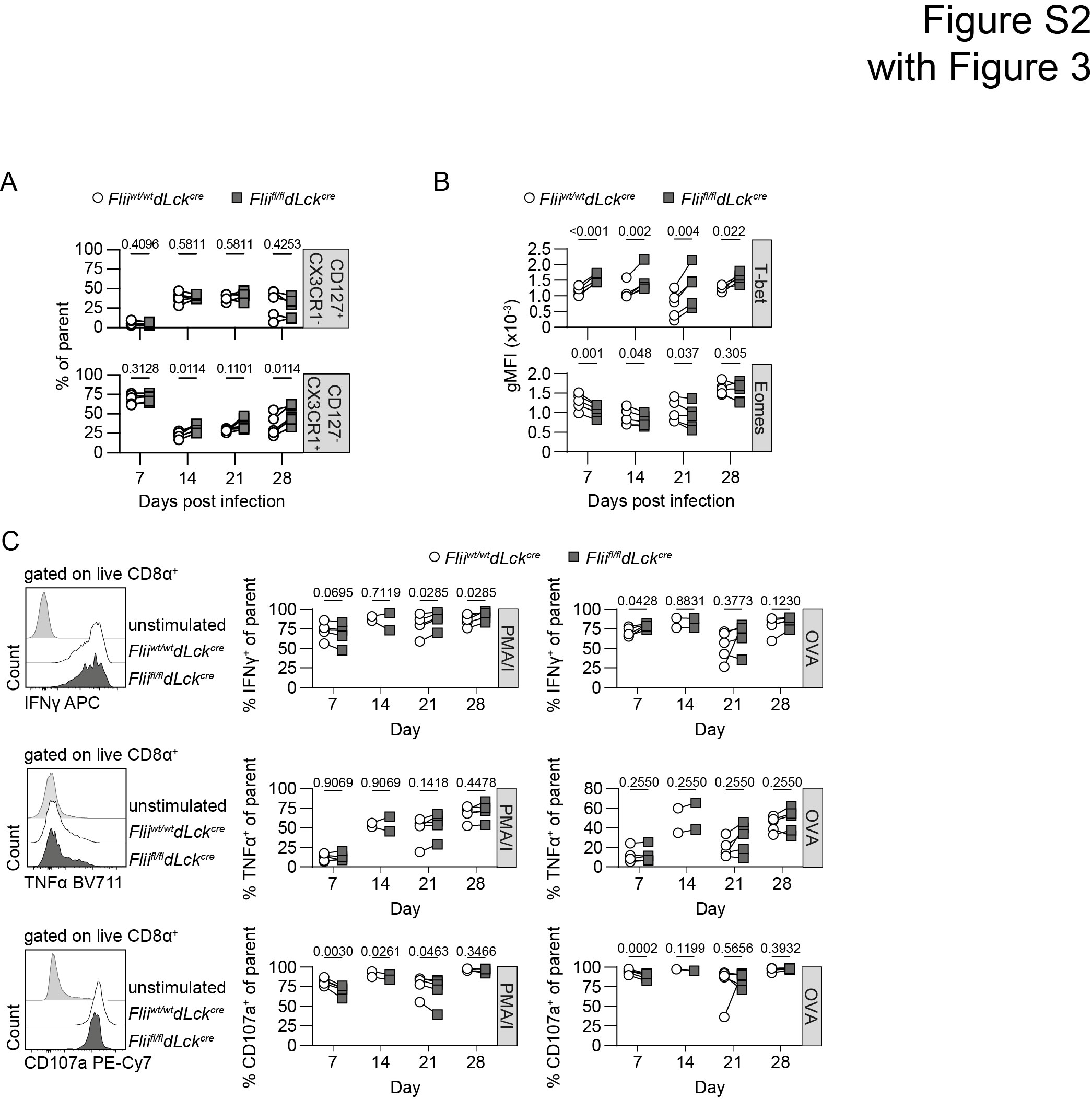

### Fig S3

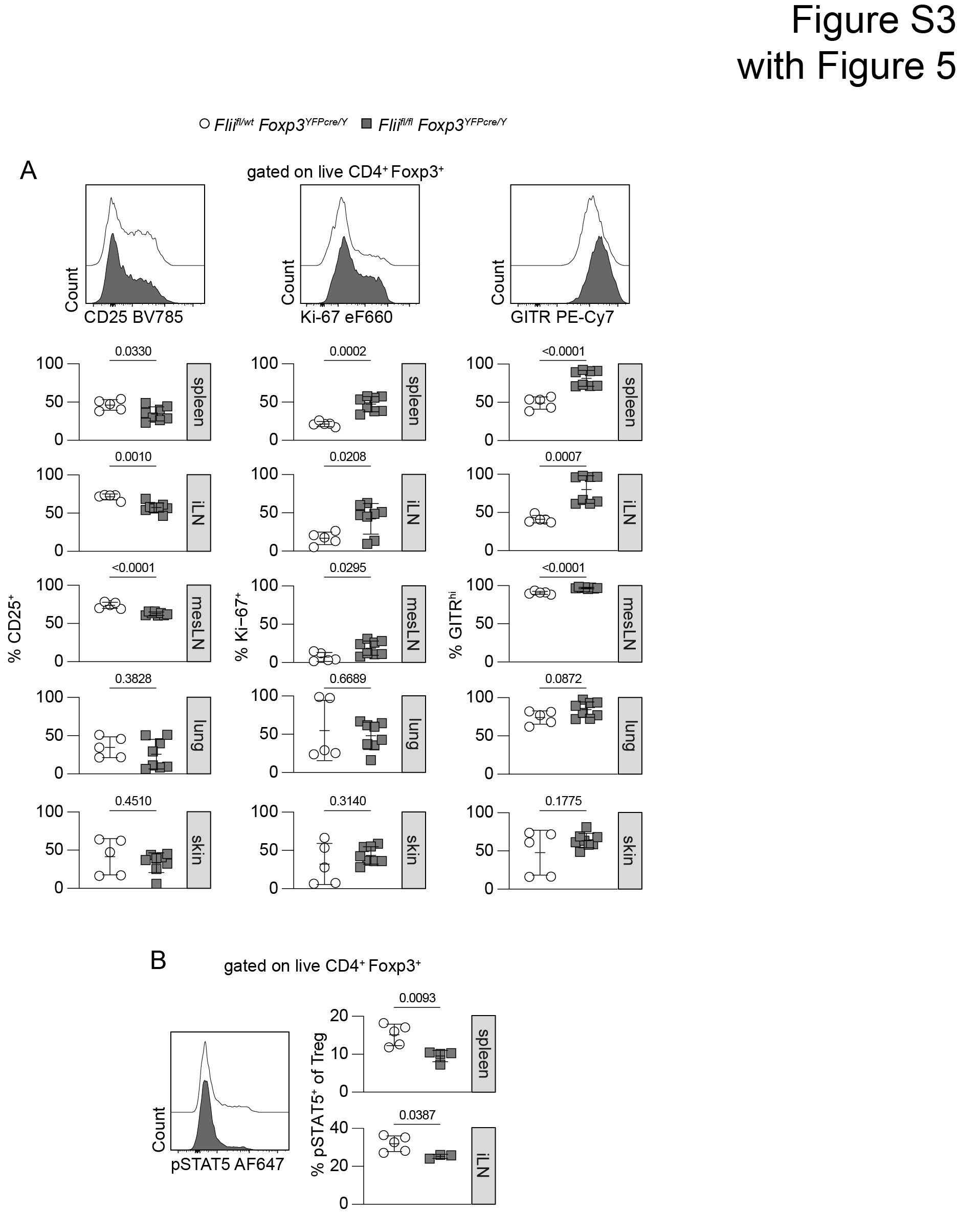

### Fig S4

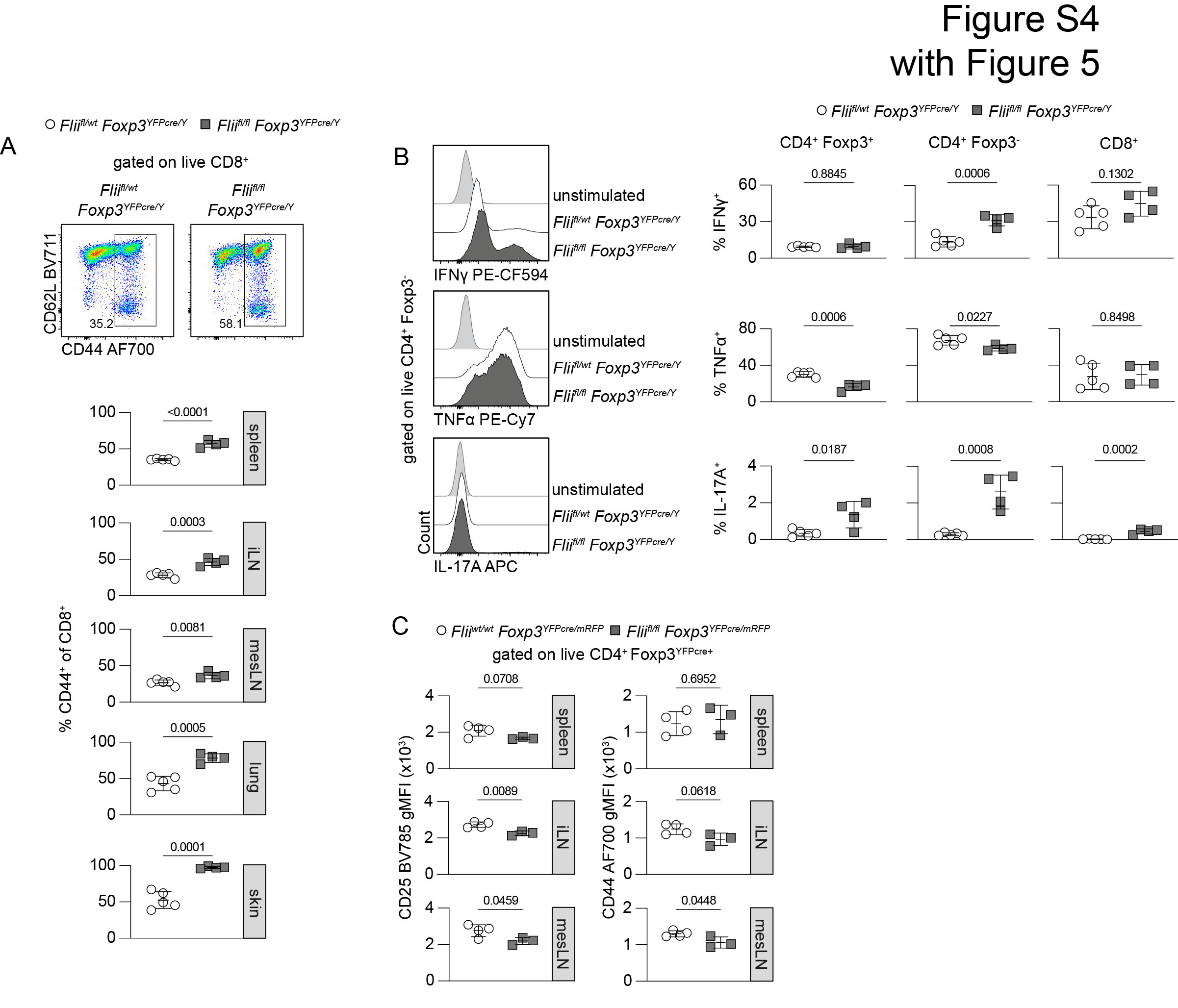
